# Multi-task learning of a deep k-nearest neighbour network for histopathological image classification and retrieval

**DOI:** 10.1101/661454

**Authors:** Tingying Peng, Melanie Boxberg, Wilko Weichert, Nassir Navab, Carsten Marr

## Abstract

Deep neural networks have achieved tremendous success in image recognition, classification and object detection. However, deep learning is often criticised for its lack of transparency and general inability to rationalize its predictions. The issue of poor model interpretability becomes critical in medical applications, as a model that is not understood and trusted by physicians is unlikely to be used in daily clinical practice. In this work, we develop a novel multi-task deep learning framework for simultaneous histopathology image classification and retrieval, leveraging on the classic concept of k-nearest neighbors to improve model interpretability. For a test image, we retrieve the most similar images from our training databases. These retrieved nearest neighbours can be used to classify the test image with a confidence score, and provide a human-interpretable explanation of our classification. Our original framework can be built on top of any existing classification network (and therefore benefit from pretrained models), by (i) adding a triplet loss function with a novel triplet sampling strategy to compare distances between samples and (ii) a Cauchy hashing loss function to accelerate neighbour searching. We evaluate our method on colorectal cancer histology slides, and show that the confidence estimates are strongly correlated with model performance. The explanations provided by nearest neighbors are intuitive and useful for expert evaluation by giving insights into understanding possible model failures, and can support clinical decision making by comparing archived images and patient records with the actual case.

## 1 Introduction

Since the overwhelming success of deep learning in the ImageNet challenge in 2012 [1], novel image recognition techniques are now based on deep learning. This is also true for histopathological image analysis, with deep learning based methods developed for mitosis detection [2], cancer classification [3], mutation prediction [4] and survival prediction [5].

Despite the breakthroughs they have made, the adoption of deep neural networks in daily clinical practice is slow. One bottleneck is that deep neural networks are often perceived as ‘black-box’ models, as it is very difficult to understand how networks make their predictions with their millions of model parameters. This issue becomes critical in computational pathology, as pathologists need to understand the rationale of a network’s decision for a certain input, if they would like to use it for diagnostic purpose. Moreover, recent studies have found deep neural networks are particularly vulnerable to adversarial examples [6]: with a small amount of permutations that are imperceptible to human, adversarial inputs can easily fool deep neural network and result in completely wrong classification, which suggest the danger of using deep neural networks without expert control.

In this paper, we aim to improve model interpretability of deep neural networks to pathologists without the need of a computational background. Inspired by the decision making process of pathologists, i.e. relating the current case to similar cases stored in their brains, we design a novel multi-task learning framework for simultaneous image classification and retrieval. In addition to cross-entropy loss used for the classification task, we add a triplet loss function to compare distance between samples [7] and a Cauchy hashing loss function to accelerate nearest neighbour search in Hamming space [8]. Through deeply retrieved nearest neighbor images, we can provide pathologists intuitive explanations of model predictions by visualizing the embedding space that is close to human perception, and also calculate an confident score by measuring the variations of the retrieved neighbours. This approach pushes classification networks in histopathology for the first time towards confident, interpretable and efficient image retrieval and hence will have a big impact on the quickly growing field of computational pathology.

## 2 Method

A schematic of our proposed multi-task learning framework for k-nearest neighbour retrieval is shown in Fig. 1. Each compartment of the framework is explained in the following subsections accordingly.

**Fig. 1.**
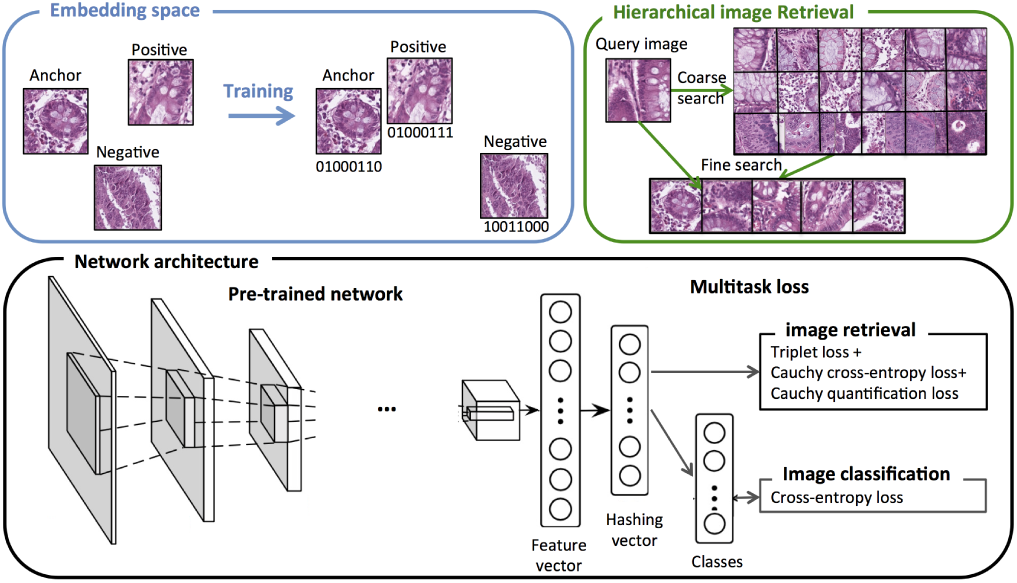
A multi-task learning framework for simultaneous image classification and retrieval. Our framework is comprised of five key components: (1) a convolutional neural network backbone for learning a deep representation of each image as a feature vector; (2) a fully-connected hash layer for transforming the feature vector into a K-bit hashing vector, which is then discretized into a binary hashing code *h*_*i*_∈{0, 1}^*K*^ by taking the sign of the each neurons in the hashing vector; (3) a triplet loss function which brings close samples from same class and push apart samples from different classes; (4) a Cauchy loss function which is a combination of Cauchy cross-entropy for similarity preserving learning and a Cauchy quantification loss for controlling the binarization error; (5) a cross-entropy loss that assesses the image classification accuracy.

### 2.1 Triplet loss with batch-hard sampling

The triplet loss has been firstly introduced by [7] for face recognition. In contrast to Siamese networks that measure pairwise distance, triplet loss considers the triangular relationship between three samples: an anchor instance *x*, a positive instance *x*^+^ that is similar to *x* (usually belonging to the same class), and a negative instance *x*^−^ that is different from *x* (usually belonging to a different class). The network is then trained to learn an embedding function *f* (.), with a loss function defined in [9]:

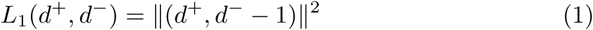

where:

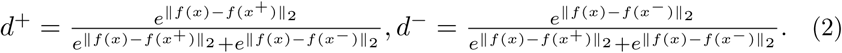

Fundamental to triplet networks is the right sampling strategy. Random sampling is usually not sufficient as most random negative images radically differ from the anchor image in the embedding space and no longer contribute to the gradients in the optimisation process. Hence, [7] proposed a batch-hard strategy which selects for each anchor sample the most distant positive (hard-positive) sample and the closest negative (hard-negative) sample. Here we propose an improved batch-hard strategy: 1) sample a balanced data set of *k* samples from each of the *n* classes; 2) compute embedding for each sample; 3) choose each sample to be an anchor, and match it with all *k −*1 positive samples; 4) for each anchor sample, choose *k* closest negative (hard-negative) samples, hence matching all anchor-positive pairs. This strategy results in *n* * *k*\* (*k−* 1) triplets when computing *n*\**k* embeddings only, which is more computational efficient than the original strategy where three embeddings were computed for one triplet. Moreover, sampling *k* hard-negative samples instead of one makes our approach more robust against outliers.

### 2.2 Cauchy loss for efficient image retrieval in Hamming space

Although the triplet network can train an efficient embedding function that preserves similarity, the resulted embedding vectors are continuous and need a *L*2 distance comparison for neighbour searching. A more efficient searching method is hashing, which compares binary codes in hamming space [10]. Recent works have focused on combining convolutional neural network with hashing methods, yielding an end-to-end framework that jointly preserves pairwise similarity and controls the quantization error [10]. Here we use the Deep Cauchy Hashing proposed in [8], which achieves superior performance over other state-of-the-art hashing approaches such as Hashnet [11]. In combination with our triplet sampling, we write the Cauchy loss function as:

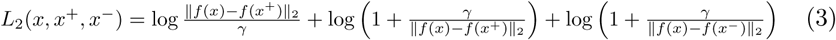

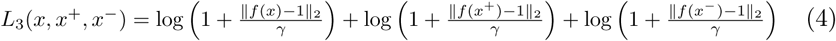

where *L*_2_ is the cross-entropy term that preserves similarities and *L*_3_ measures the quantification error before and after discretization, where we generate a binary hashing code by taking the sign of each neuron of the hashing vector. The scale parameter *γ* controls the decaying speed of the probability of the Cauchy distribution: a smaller *γ* will impose more force to concentrate similar samples into a small Hamming radius. Here we choose *γ* to be *K/*2, where *K* is the bit number of our hashing code.

### 2.3 Cross-entropy loss as an auxiliary classification task

To enable our framework for classification, we add a classification layer after the hashing layer, in which a cross-entropy loss is used to minimize the discrepancy between prediction and ground-truth labels. The addition of a classification function also allows us to compare the performance of our framework to previous work of [5], which use standard classification networks.

### 2.4 Hierarchical image retrieval

In the testing phase, for each query image we adopt a coarse-to-fine search strategy for rapid and accurate image retrieval. We first retrieve a candidate pool with similar binary hashing codes after discretization within a small Hamming radius (e.g. of 1) from the query image. To further filter the images with similar appearance, we extract the feature vector (one layer before the hashing vector) and rank the retrieved samples (see Fig. 1) using a *L*2 distance of the feature vector. In our implementation, we use the built-in functions *BallTree* and *cKDTree* of the scikit-learn toolbox for nearest neighbour searching in Hamming space and Euclidean space. As one important purpose of our image retrieval is for expert evaluation, we limit the number of retrieved images for each query image to be 10.

### 2.5 Confidence measurement

The retrieved nearest neighbours of a given query image also provide a straight-forward confidence measure of our prediction on that image by simply counting the frequency of the predicted class in the retrieved neighbourhood.

## 3 Results and Discussions

### 3.1 Experimental data

In order to evaluate our framework we use the colorectal cancer (CRC) histology dataset [5]. It contains more than 100,000 hematoxylin-eosin (HE)-stained image patches from 86 CRC tissue slides from the NCT biobank and the UMM pathology archive (NCT-CRC-HE-100K) and a testing data set of 7,180 image patches from 25 CRC patients from an independent cohort (CRC-VAL-HE-7K). Both datasets are created by pathologists by manually delineating tissue regions in whole slide images into the following nine tissue classes: adipose tissue, background, cellular debris (comedonecrosis), lymphocytes, extra-cellular mucus, smooth muscle (lamina muscularis mucosae), normal colon mucosa, cancer-associated stroma, and neoplastic cell population (CRC epithelium). CRC epithelium was exclusively derived from human CRC specimen (primary and metastatic). Normal tissue such as smooth muscle and adipose tissue was mostly derived from CRC surgical specimen, but also from upper gastrointestinal tract specimen (including smooth muscle from gastrectomy) in order to maximize variability in this training set. The created non-overlapping image patches are 224 × 224*px* (112 × 112*µm*) and have a approximately equal distribution among the nine tissue classes. [5] trained a classification network on NCT-CRC-HE-100K and reach 98.8% accuracy on the test split of the dataset and 94.3% accuracy on the independent test set (CRC-VAL-HE-7K).

### 3.2 Evaluation of image classification

To train our framework we split the training data (NCT-CRC-HE-100K) into 70% training set,15% validation set and 15% test set. The independent cohort (CRC-VAL-HE-7K) is used for testing purpose only. We choose convolutional neural networks of different architectures and replace the last layer of each network with our hashing and classification layers (see Sec. 2 and Fig. 1). To train each network, we initiate it with ImageNet pretrained weights, train our added layers first and then fine tuning the entire network. In additional to different network architectures, we also examine the influence of multi-task learning by comparing the classification performance when training with multi-task loss vs. training with cross-entropy loss for classification only. The classification accuracy we achieve is comparable to the results reported in [5], suggesting our networks are properly trained (see Table 1). Moreover, we demonstrate that the multi-task learning improves the classification performance on an unseen test set, suggesting the advantage of using our multi-task loss combination. It also illustrates that there is a domain shift between the histology images from the two different cohorts, so the network that achieves the best performance on the internal test set of NCT-CRC-HE-100K does not generalize best on the indepedent CRC-VAL-HE-7K test set.

**Table 1.** Evaluation of classification accuracy on both test set of NCT-CRC-HE-100K and an independent test set of CRC-VAL-HE-7K. *The results of VGG19 is directly quoted from [5] and are shown here as a comparison.

| Testing accuracy | Multitask Resnet18 | Resnet18 | Resnet34 | Resnet50 | VGG19* |
| --- | --- | --- | --- | --- | --- |
| NCT-CRC-HE-100K (%) | 98.6 | 98.5 | 98.8 | **99.4** | 98.8 |
| CRC-VAL-HE-7K (%) | **95.0** | 94.4 | 94.2 | 93.6 | 94.3 |

### 3.3 Evaluation of image retrieval

To evaluate image retrieval, we use the entire NCT-CRC-HE-100K set as our database and the independent CRC-VAL-HE-7K set as query images. As explained in Sec. 2.4, for a query image we use the coarse-to-fine strategy to retrieve its nearest neighbours. To make a comparison, we formulate a baseline neighbour searching method using classification: we amend our coarse search to compare hashing vectors without discretization using *L*_2_ distance and to retrieve 100 neighbours as the candidate pool for the next fine search. We measure our retrieval precision for each query image by counting the number of true neighbours, i.e. belonging to the same class, among the top 10 retrieved samples, as proposed in [8][11]. Over 6000 images out of our 7180 query images reach a perfect retrieval precision of 10 true neighbours by using our multi-task network (Fig. 2), which is around 30% higher than that achieved by the baseline classification network (4697 images). This suggests that the embedding space created by multitask framework is more compact, i.e. a sample is surrounded predominantly by neighbours of its own class. By contrast, in the embedding space created by baseline classification, a sample is more mixed with neighbours of different classes. A dispersed embedding could be one reason that classification networks are vulnerable to attacks of adversarial samples [6].

**Fig. 2.**
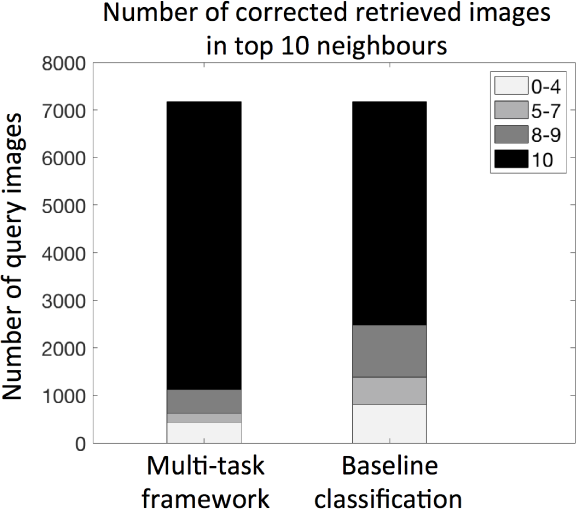
Multi-task learning retrieves more correct images than simple classification.

Fig. 4 shows exemplary results of our image retrieval. The first query image is a patch of cancer-associated stroma. While the classification network confuses it with patches of smooth muscle in healthy tissue due to their similar color appearance, our multi-task network, by contrast, is not fooled by the colour variations and is able to reach perfect retrieval. The second query image is a patch of colorectal adenocarcinoma epithelium, which is mixed with normal colon mucosa by the classification network but not by the multi-task network.

**Fig. 3.**
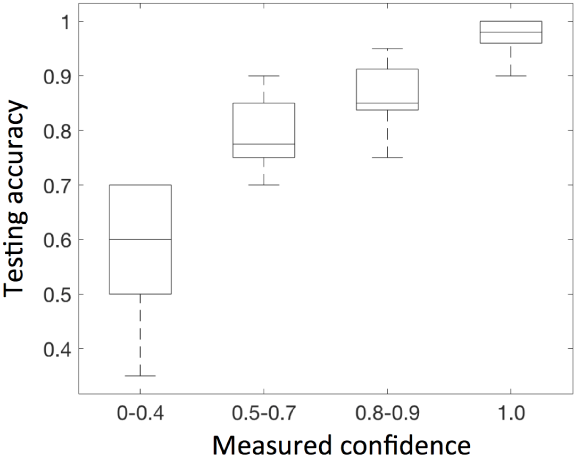
Our confidence measure is highly correlated with their testing accuracy (variations come from multiple testing of batches of size 50).

**Fig. 4.**
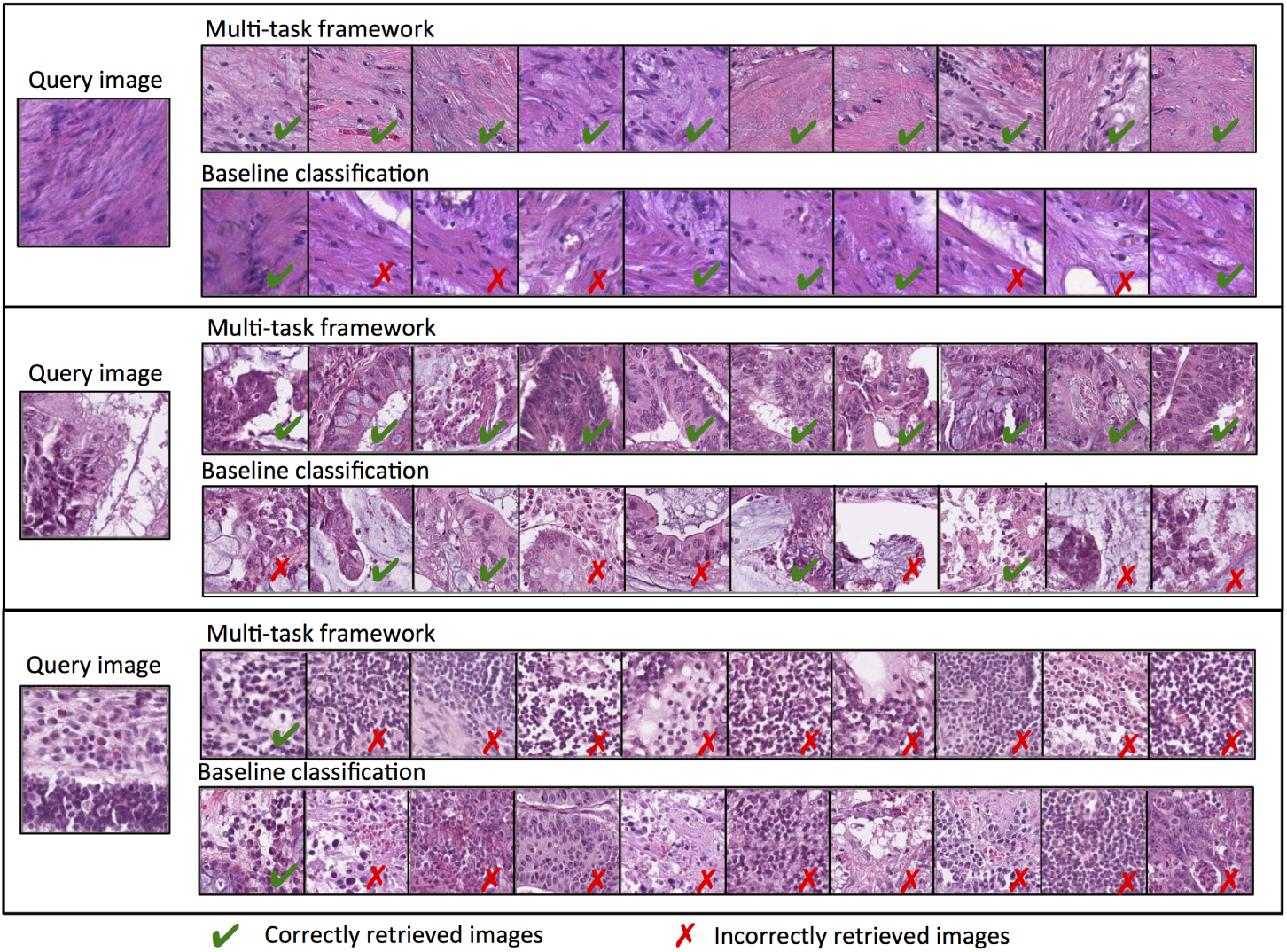
The top 10 retrieved images returned by our multi-task framework as compared to a baseline classification network for exemplary cases. See text for explanation.

Fig. 3 shows that the confidence measurement of our framework is highly correlated with the actual performance of our classification on the testing set. One exemplary low confident retrieval case captured by our framework is shown in the last row of Fig. 4, the query image is annotated as normal colon mucosa yet is considered to be mostly lymphocytes, debris, cancer-associated stroma and colorectal adenocarcinoma epithelium by our framework. An expert pathologist also reviewed the case and did not agree with its original annotation as normal colon mucosa, though a more definite conclusion could not be reach due to the limited context provided by this patch. Our framework can be used to highlight these uncertain cases for review by more than one pathologists.

### 4 Conclusion

We propose a novel multi-task learning framework for simultaneous image classification and retrieval. Our objective function is composed of a triplet loss function to compare distance between samples, a Cauchy hashing loss function to accelerate nearest neighbour search in Hamming space and a classic cross-entropy loss to assess classification performance. We demonstrate that such a multi-task learning framework learns a more compact and accurate embedding space as compared to classic classification networks and allows medical experts to explore and check the embedding space without the need of in-depth machine learning knowledge. Moreover, we illustrate that the confidence measure provided by the variations of the retrieved neighbourhood is highly correlated with the model performance and hence can be used to select low confident predictions for expert review. Our framework can be turned into a very useful tool to support clinical decision making of pathologists by comparing archived images and patient records with the actual case.

## References

1. Krizhevsky, A. et al: ImageNet Classification with Deep Convolutional Neural Networks, NIPS (2012) 1097–1105.

2. Ciresan, DC et al: Mitosis detection in breast cancer histology images with deep neural networks., MICCAI (2013) 411–8.

3. Esteva, A et al: Dermatologist-level classification of skin cancer with deep neural networks., Nature (2017) 542 (7639) 115–118.

4. Coudray, N et al: Classification and mutation prediction from nonsmall cell lung cancer histopathology images using deep learning., Nature Medicine (2018) 24 (10) 1559–1567.

5. Kather, JN et al: Predicting survival from colorectal cancer histology slides using deep learning: A retrospective multicenter study., PLOS Medicine (2019) 16 (1) e10027300.

6. Paschali, M et al: Generalizability vs. Robustness: Investigating Medical Imaging Networks Using Adversarial Examples., MICCAI (2018) 493–501.

7. Shroff, F et al: FaceNet: A unified embedding for face recognition and clustering., CVPR (2015) 815–823.

8. Cao, Y et al: Deep Cauchy Hashing for Hamming Space Retrieval., CVPR (2018) 1229–1237.

9. Hoffer, E et al: Deep Metric Learning Using Triplet Network., ICLR (2015) 84–92.

10. Wang, J et al: A Survey on Learning to Hash., IEEE Transactions on Pattern Analysis and Machine Intelligence (2018) 40(4) 769–790.

11. Cao, Z et al: HashNet: Deep Learning to Hash by Continuation., ICCV (2017) 5609–5618.

